# gauseR: Simple methods for fitting Lotka-Volterra models describing Gause’s “Struggle for Existence”

**DOI:** 10.1101/2020.03.16.993642

**Authors:** Lina K. Mühlbauer, Maximilienne Schulze, W. Stanley Harpole, Adam T. Clark

## Abstract

The ecological models of Alfred J. Lotka and Vito Volterra have had an enormous impact on ecology over the past century. Some of the earliest – and clearest – experimental tests of these models were famously conducted by Georgy Gause in the 1930’s. Although well known, the data from these experiments are not widely available, and are often difficult to analyze using standard statistical and computational tools. Here, we introduce the `gauseR` package, a collection of tools for fitting Lotka-Volterra models to time series data of one or more species. The package includes several methods for parameter estimation and optimization, and includes 42 datasets from Gause’s species interaction experiments and related work. Additionally, we include with this paper a short blog post discussing the historical importance of these data and models, and an R vignette with a walk-through introducing the package methods. The package is available for download at github.com/adamtclark/gauseR. To demonstrate the package, we apply it to several classic experimental studies from Gause, as well as two other well-known datasets on multi-trophic dynamics on Isle Royale, and in spatially structured mite populations. In almost all cases, models fit observations closely, and fitted parameter values make ecological sense. Taken together, we hope that the methods, data, and analyses that we present here provide a simple and user-friendly way to interact with complex ecological data. We are optimistic that these methods will be especially useful to students and educators who are studying ecological dynamics, as well as researchers who would like a fast tool for basic analyses.

## Introduction

A century ago, Alfred J. Lotka (1920, 1925) and Vito Volterra (1926) published their canonical works concerning the mathematics of species interactions. These theories were among the first to suggest that the dynamics of complex ecological systems could be described through simple underlying equations (Giorgio 1988). Inspired by the possibility of understanding natural systems through mathematics, Georgy Gause stated in his “Struggle for Existence” (1934b):

> *“[…] in order to penetrate deeper into the nature of these phenomena we must combine the experimental method with the mathematical theory, a possibility which has been created by the brilliant researches of Lotka and Volterra.”*

To match the strong assumptions and narrow scope of Lotka and Volterra’s basic equations, Gause constructed simple experimental microcosms, in which he was able to show that interactions between pairs of species influenced their abundances and dynamics in ways that accorded with theoretical predictions (Gause 1934a,b). These experiments laid the groundwork for a century of experiments based on mathematics as a conceptual framework. It has since become a primary goal of ecology to explain empirical observations with mathematical theory.

Despite their simplicity, the Lotka-Volterra equations are still the primary models used to describe species interactions. At their most basic, these models describe the changes per unit time in the abundance of a species, *N*_*i*_, as

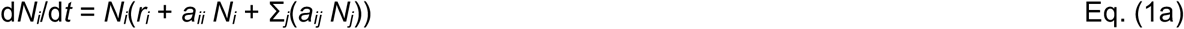

where *r*_*i*_ is the intrinsic growth rate, which describes the growth of a species at low density and in the absence of other species, *a*_*ii*_ describes the effect of species *i* on its own growth, and *a*_*ij*_ describes the effect of species *j* on the growth of species *i*. The form that we use here differs somewhat from that often portrayed in some textbooks, as we leave *r*_*i*_ inside the parenthesis and therefore do not standardize interaction coefficients by the growth rate (thus, we signify these as *a* rather than *α*). If we divide both sides of Eq. (1a) by *N*_*i*_ to calculate the “per-capita” growth rate of species *i*, the model can be written as a simple linear equation

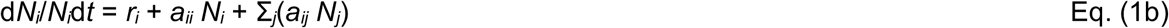

This form is particularly useful for analyzing empirical data, as the parameters can be estimated using ordinary least squares regression of per-capita growth rates against species abundances, where *r*_*i*_ is the y-intercept intercept, *a*_*ii*_ is the slope with respect to species *i*’s own abundance, and *a*_*ij*_ is the slope with respect to other species (Lehman *et al*. 2020). Note that in a single species, this relationship can also be expressed in terms of the carrying capacity, *K*_*i*_, which describes the equilibrium abundance reached by a species growing in the absence of other species, and falls on the x-intercept (Fig. 1). For those who are less familiar with regression or dynamical systems modelling, we have included a more in-depth discussion of this approach in the vignette accompanying our R package. We can solve for *K*_*i*_ in terms of *a*_*ii*_ and *r*_*i*_ by setting Eq. (1b) equal to zero, and solving for *N*_*i*_, which yields

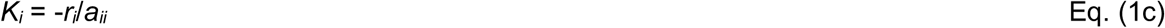

**Figure 1:**
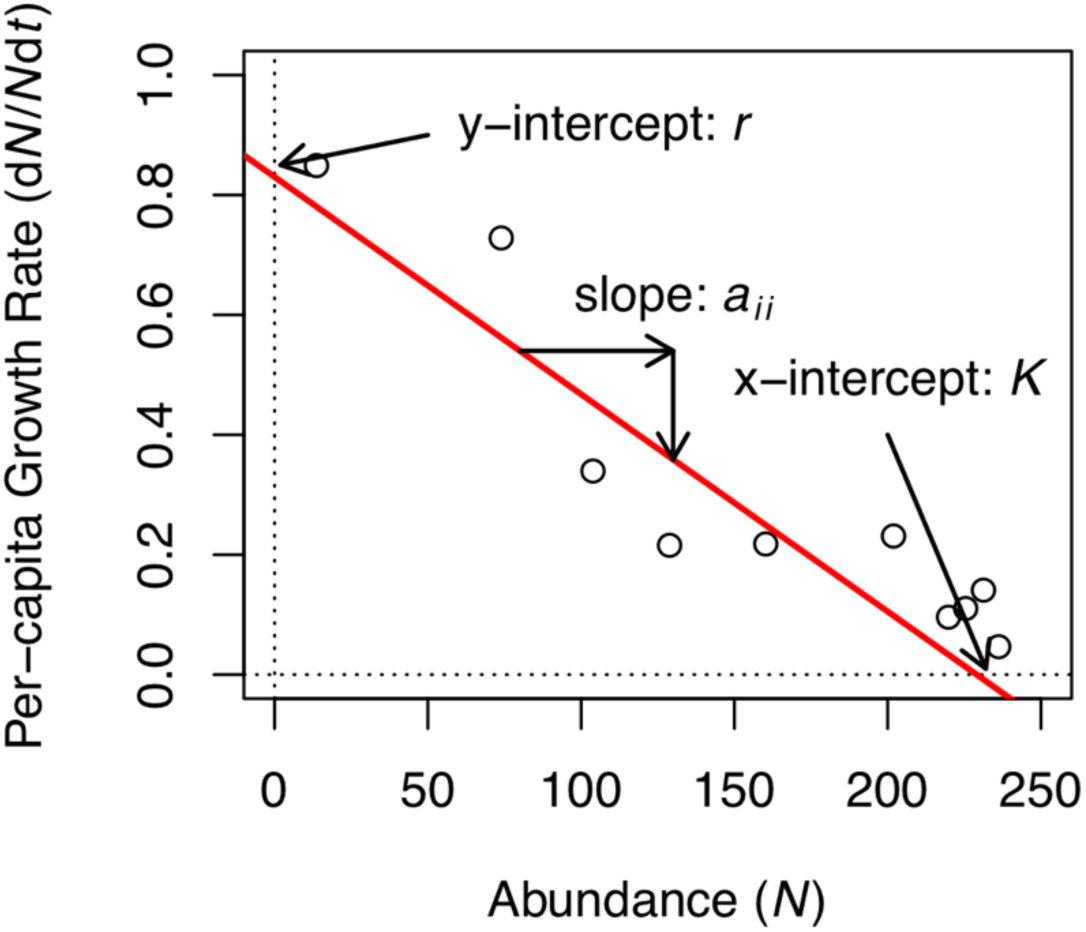
Figure comparing per-capita growth rate (following Eq. 1b) vs. abundance for *Paramecium aurelia* grown in monoculture. Note that when plotted in this manner, the parameter values for the Lotka-Volterra equations can be easily “read” off the graph, which *r* as the y-intercept, *K* as the x-intercept, and *a*_*ii*_ as the slope. In multi-species systems, interaction coefficients *a*_*ij*_ (i.e. effect of species *j* on species *i*) can be similarly computed based on the slope of d*N*_*i*_/*N*_*i*_d*t* vs. *N*_*j*_.

Because the Lotka-Volterra models make no specific assumptions about the mechanisms underlying species interactions, they can be parameterized in ways that approximate any combination of underlying mechanisms, at least locally around equilibrium (MacArthur 1970; Chesson 2000; Letten & Stouffer 2019). For example, in a system with two species, if both *a*_*ij*_ and *a*_*ji*_ are less than zero, species suppress each other’s growth, indicating competition. If both parameters are positive, then both species benefit from the other’s presence, indicating mutualism. Alternatively, if *a*_*ij*_ < 0 and *a*_*ji*_ > 0, then species *i* is suppressed by species *j* whereas species *j* benefits from the presence of species *i*, indicating a predator-prey relationship.

Gause’s original experimental work still provides some of the best examples of these different kinds of interspecific interactions. However, because of the limited analytical tools available to him in the early twentieth century, Gause was only able to fit models to a small subset of his experimental data. Moreover, because of the journal formatting conventions of his time, much of Gause’s data was never officially published. Thus, although they are widely used as teaching tools and widely cited as canonical examples of competitive and predator-prey interactions, very few of his results have ever been quantitatively analyzed using the Lotka-Volterra equations.

Here, we introduce the `gauseR` package, which includes a collection of functions for confronting empirical observations with the classical Lotka-Volterra interaction models. The package combines methods for data formatting, model simulation, and parameter estimation into a simple-to-use, automated wrapper function. The package also acts as a digital repository for Gause’s experiments on the “Struggle for Existence”, and includes all available data from his 1934 book and associated publications. Below, we introduce the functions and basic workflow for the package, and explain the underlying theory. To demonstrate the package’s functionality, we analyze two well-known datasets from Gause (1934a), as well as classic examples of multi-species interactions from McLaren & Peterson (1994) and Huffaker *et al*. (1963). Additionally, we discuss some limitations and possible extensions of the optimization methods used in the package. We foresee that `gauseR` will be especially helpful for researchers who seek simple tools for analyzing their data, as well as a teaching aid for introducing students to the Lotka-Volterra equations and to Gause’s experiments.

## Methods

### Functions and workflow

The general purpose of the functions in the `gauseR` package is to fit the general Lotka-Volterra model in Eqs. (1a-b) to observational time series data of one or more interacting species. The package, along with a walk-through vignette, is available at github.com/adamtclark/gauseR. Functions are described in detail in Table 1. In general, this procedure will require estimating values for three types of parameters: starting abundances (*N*_*0i*_), intrinsic growth rates (*r*_*i*_) and interaction coefficients (*a*_*ij*_).

**Table 1:** Brief descriptions of the `gauseR` functions.

| Function | Description |
| --- | --- |
| **get_lag** | Calculates time-lagged observations for variable x, separated by treatment. |
| **percap_growth** | Calculates per-capita growth rate, using log ratios. |
| **get_logistic** | Calculates logistic growth for a population. |
| **lv_interaction** | Calculates $dn/dt$ for n species in a Lotka-Volterra system. |
| **lv_interaction_log** | Calculates $dn/dt$ for n species in a Lotka-Volterra system, with log-transformed abundances. This form can be helpful for optimization routines where species abundances are close to zero. |
| **lv_optim** | Identifies optimal parameter values for a Lotka-Volterra interaction system by simulating dynamics using differential equations. |
| **gause_wrapper** | Automatically runs routine for finding starting values and optimal parameter values for a Lotka-Volterra interaction system. |
| **test_goodness_of_fit** | Generates an $R^2$-like goodness of fit index, based on the squared difference between observed and predicted values. Index values of 1 indicate a very close fit, whereas values near or below zero indicate a poor fit. |
| **ode_prediction** | Calculates expected abundances for all n species in a community. This function is potentially useful in combination with other optimizer software, e.g. as might be used for hypothesis testing. |

For a single species system, dynamics can be directly fit to the Logistic growth model using the `get_logistic` function. For systems with two or more species, fitting proceeds in four steps. First, time-lagged abundances for each species are calculated using the `get_lag` function, yielding pairs of abundance values separated by a fixed time interval for each species. Second, these time-lagged abundances are used to estimate per-capita growth rates at different time points, using the `percap_growth` function. This function approximates dynamics following the linearized equation

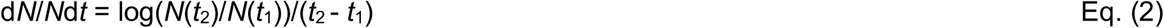

Third, linear regressions are used to test how per-capita growth rates vary as a function of conspecific and heterospecific species abundances, from which the parameters in Eq. (1b) can be estimated. Finally, because parameter estimates from linearized growth rates often do not match observations for long or complex time series, the `lv_interaction` and `lv_optim` function are used to directly simulate dynamics and tune parameters to match model outputs to observations, which can greatly improve model performance (Carrara *et al.* 2015; Rosenbaum & Rall 2018; Maynard *et al.* 2019). All four of these steps are automated in the `gause_wrapper` function, which takes observed species abundances as an input, and returns fitted parameter estimates. We suggest this option for most users.

To facilitate optimization, we include two additional functions. First, the `lv_interaction_log` function simulates dynamics for log-transformed species abundances. This approach is helpful in systems where abundances approach zero, which can cause integration issues (e.g. predictions of negative abundances). This function is the default for `gause_wrapper`. Lastly, the `ode_prediction` function simulates dynamics for *n* species given a set of input parameter values, and returns estimates of species abundances as a single column vector. This function is potentially useful for users who wish to apply other, more sophisticated optimization routines.

### Data and example analyses

The `gauseR` package includes a wide range of digitized data from Gauses’ experimental work, as well as some other well-known multi-species time series. Full metadata and citations for these datasets are available in the help documentation for the package. Dataset names refer to the source and figure or table from which the data are drawn. Whenever possible, data were taken from tables in Gause’s publications or appendices. Otherwise data were digitized from figures using WebPlotDigitizer (Rohatgi 2015).

To demonstrate how to apply our functions, we also conduct example analyses using four of these datasets. First, `gause_1934_science_f02_03` shows Gause’s classic competition experiment between *Paramecium aurelia* and *P. caudatum*, including both species grown in monoculture and in mixture (Gause 1934a,b). We use these data to demonstrate parameter fitting methods for logistic growth and Lotka-Volterra competition. Full source code for these analyses are available in the help documentation for the `get_logistic` and `lv_interaction` functions. Second, `gause_1934_science_f01` includes interactions between the prey species *P. caudatum* and predator species *Didinium nasutum*, with which we demonstrate the difference between estimates derived from linearized models vs. from the optimizer fit to the full simulated dynamics. These analyses are shown in the help documentation for the `lv_optim` function. Third, to demonstrate parameter fitting in a system with more than two species, we use `mclaren_1994_f03`, which summarizes dynamics of wolves (*Canis lupus*), moose (*Alces alces*), and fir trees (*Abies balsamea*) on Isle Royale from McLaren & Peterson (1994). Finally, to demonstrate a case where classic Lotka-Volterra interaction models are unable to match observations, we analyze dynamics of the prey species *Eotetranychus sexmaculatus* and predator species *Typhlodromus occidentalis* from Huffaker’s classic mite experiments (Huffaker *et al*. 1963). Data are available in `huffaker_1963`. Source code for the analyses of the McLaren and Huffaker datasets are available in the supplement.

## Results

In most cases, the simple Lotka-Volterra model in Eqs. (1a-b) accurately describe observed dynamics, and provide interpretable parameter values. For both logistic growth (Fig. 2) and competitive interactions between the two *Paramecium* species (Fig. 3), observations closely matched model predictions. Fitted parameter values suggest that both species are subject to self-limitation in monoculture (i.e. *a*_*ii*_ and *a*_*jj*_ < 0), and indicate competitive interactions in mixture (i.e. *a*_*ij*_ and *a*_*ji*_ < 0) (Table 2a-b).

**Table 2:**
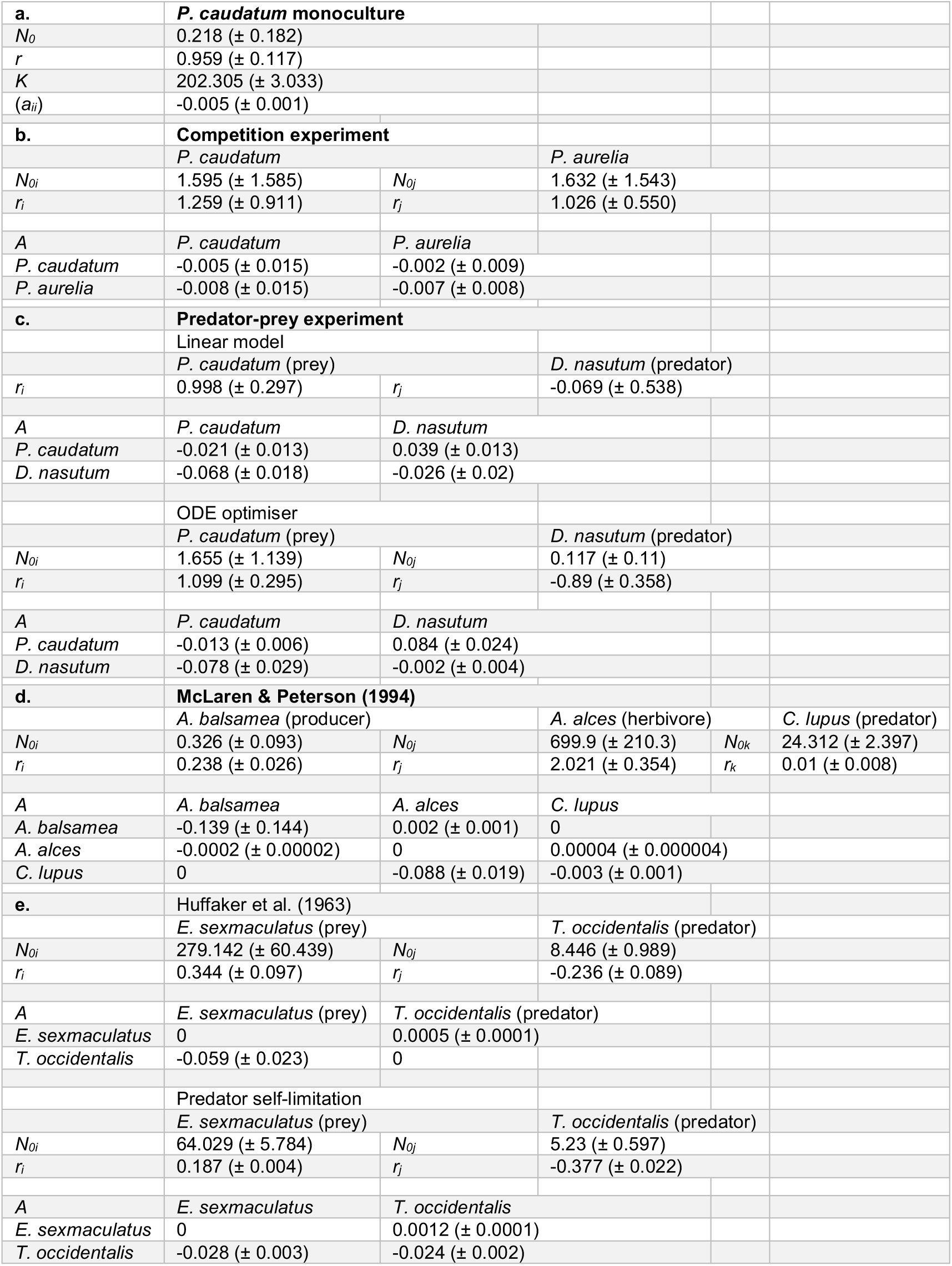
Fitted parameter values for each analysis described in the main text. Values show parameter estimates ± one standard error. For interaction matrices *A*, parameters show the effect of the species listed in the *row* on the species listed in the *column* (i.e. *a*_*ij*_ is listed in row *j* and column *i*). Note that standard errors are approximated from the numerically differentiated Hessian values calculated for the log-transformed parameters, based on the difference between the mean estimate and the lower standard deviation in log space, and are therefore only meant as rough approximations of the degree of uncertainty.

**Figure 2:**
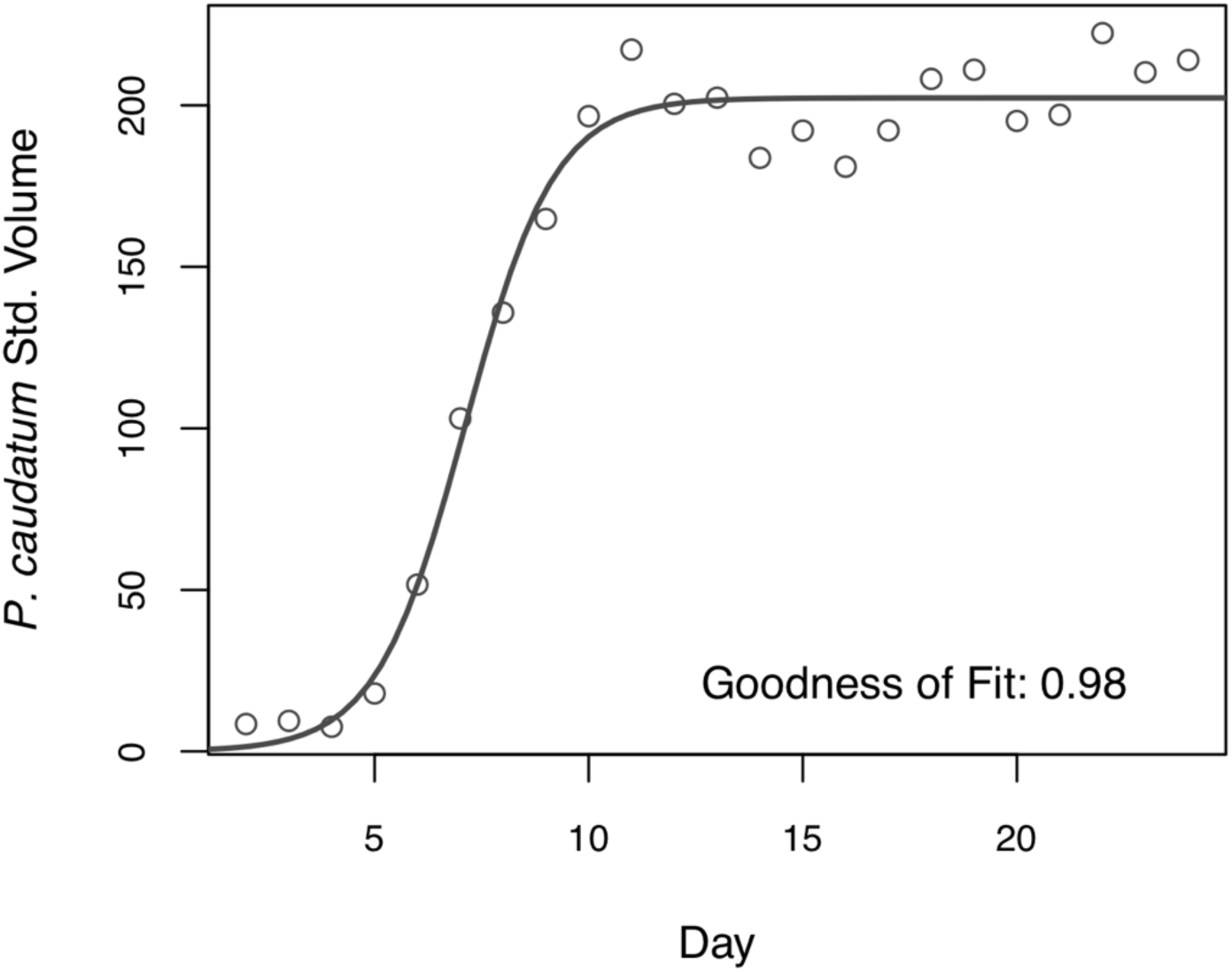
Logistic growth for size standardized volume of *Paramecium caudatum* grown in monoculture on Osterhout medium over 24 days, from Gause (1934a). Points show observations, and line shows fitted logistic growth curve, calculated with the `get_log()` function, that can be used to fit the logistic growth function to time series of an individual species. Goodness of fit shows results from the `tests_goodness_of_fit()` function. Recall that values near 1 imply a close correspondence between observations and predictions. See Table 1a for parameter values.

**Figure 3:**
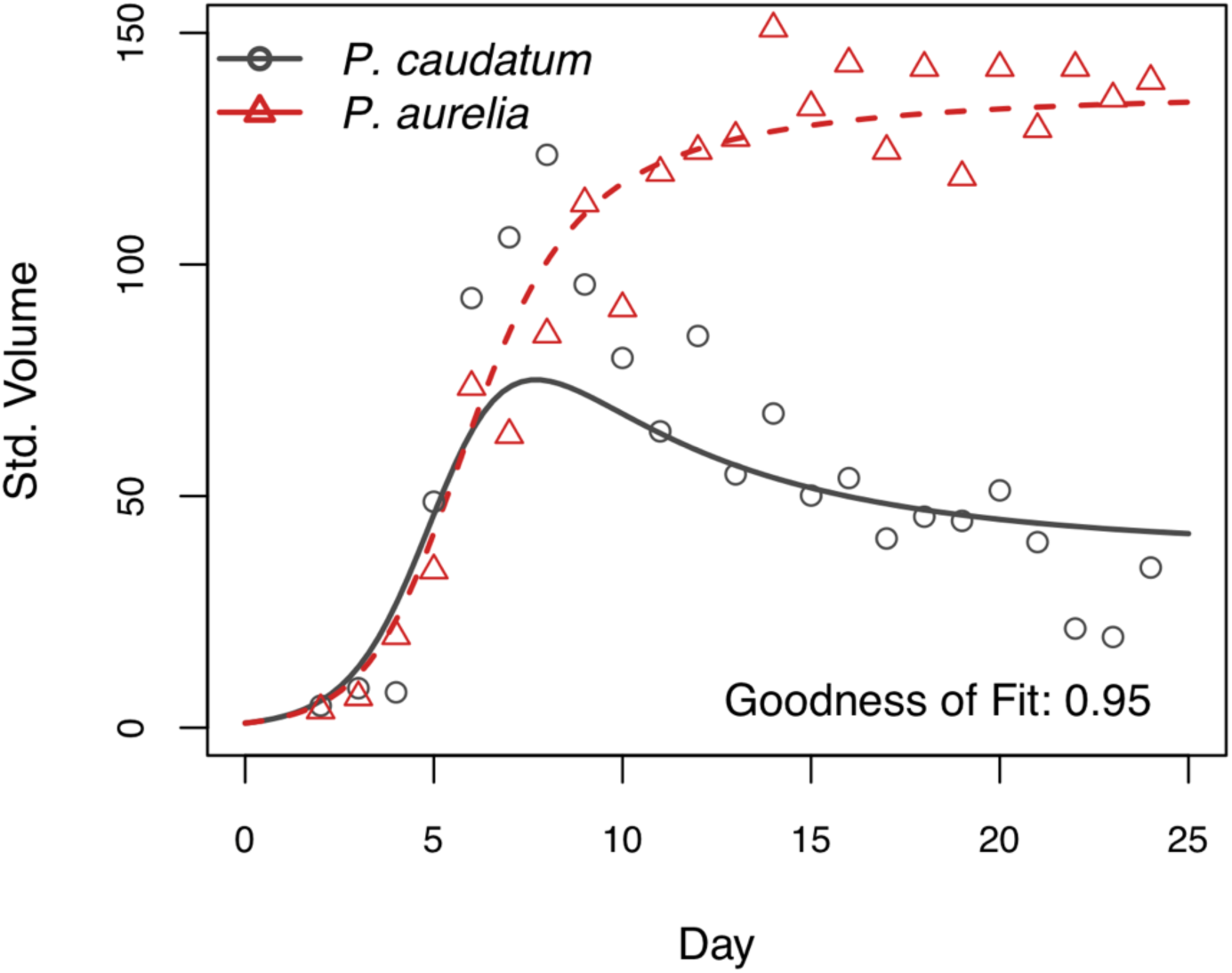
Lotka-Volterra competition for size standardized volume of *Paramecium caudatum* and *Paramecium aurelia* grown in mixed population over 24 days, from Gause (1934a). Points show observations, and lines show growth curves fitted using the `lv_interaction()` function, which can be used to simulate dynamics in multi-species mixtures following the classic Lotka-Volterra equations. See Table 1b for parameter values.

For predator-prey interactions between *D. nasutum* and *P. caudatum*, we found that parameter estimates from regressions fitted to the linearized growth rates failed to match observed dynamics (Fig. 4; Table 2c). However, predictions improved substantially after using the optimizer to tune parameters. For the wolf-moose-fir system of McLaren *et al*. (1994), results from the optimizer also matched observed dynamics (Fig. 5). Because we expected no direct interactions between wolves and fir trees, we set these interactions to zero prior to analyses. Additionally, because McLaren *et al.* hypothesized that moose populations are subject to top-down control in this system, we set self-limitation for moose to zero (Table 2d).

**Figure 4:**
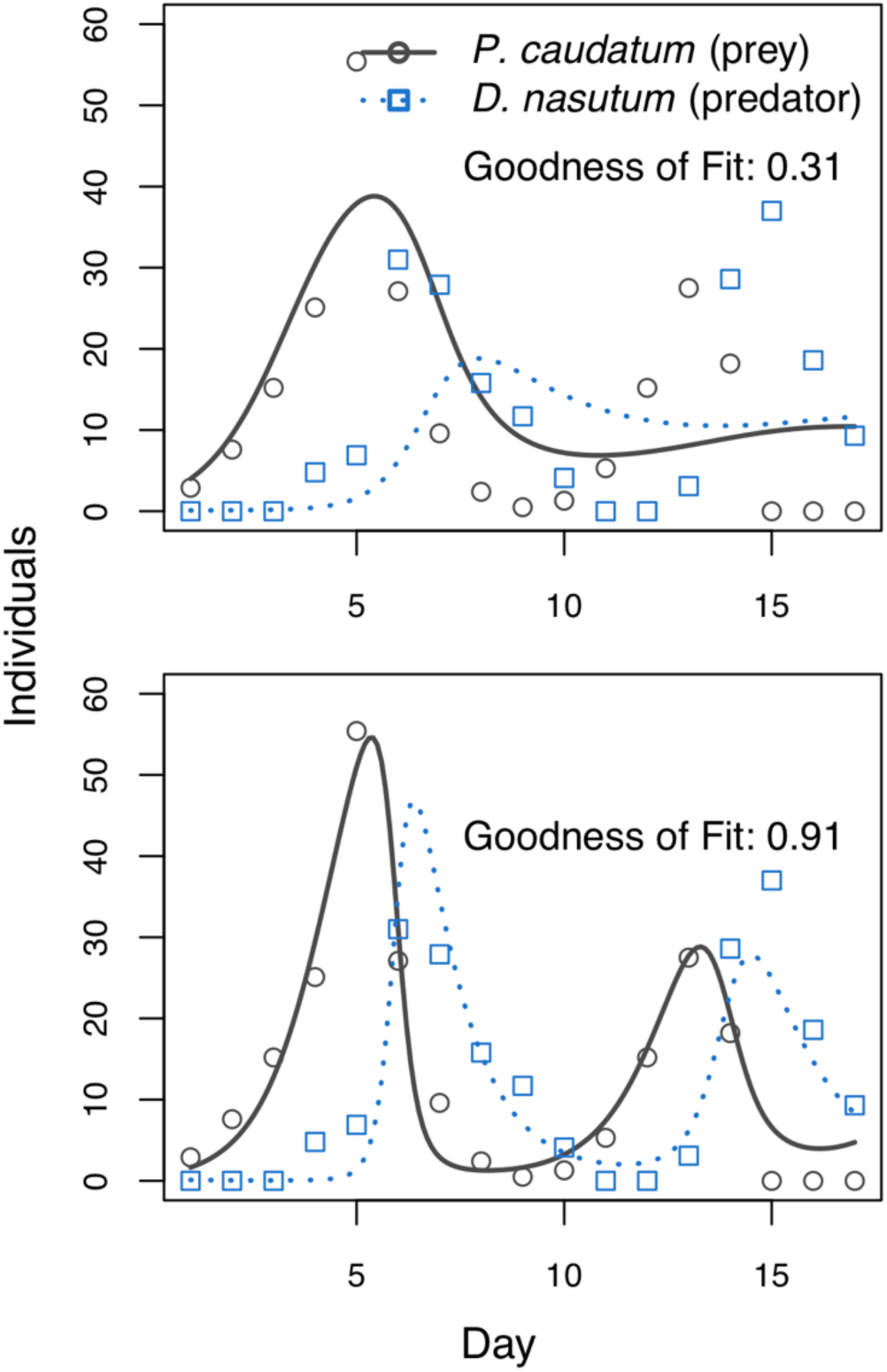
Predator-prey interactions between *Didinium nasutum* and *Paramecium caudatum* grown in mixture over 17 days, from Gause (1934a). Additional individuals of both species were added to the mixture periodically to prevent local extinction. Points show observations, and lines show fitted growth curves. Top panel shows results for model fitted using linear regressions of species abundances vs. per-capita growth rate, whereas bottom panel shows results for parameters fitted using the simulated differential equations using the `lv_optim()` function. Note that goodness of fit is varies substantially between the two methods. See Table 1c for parameter values.

**Figure 5:**
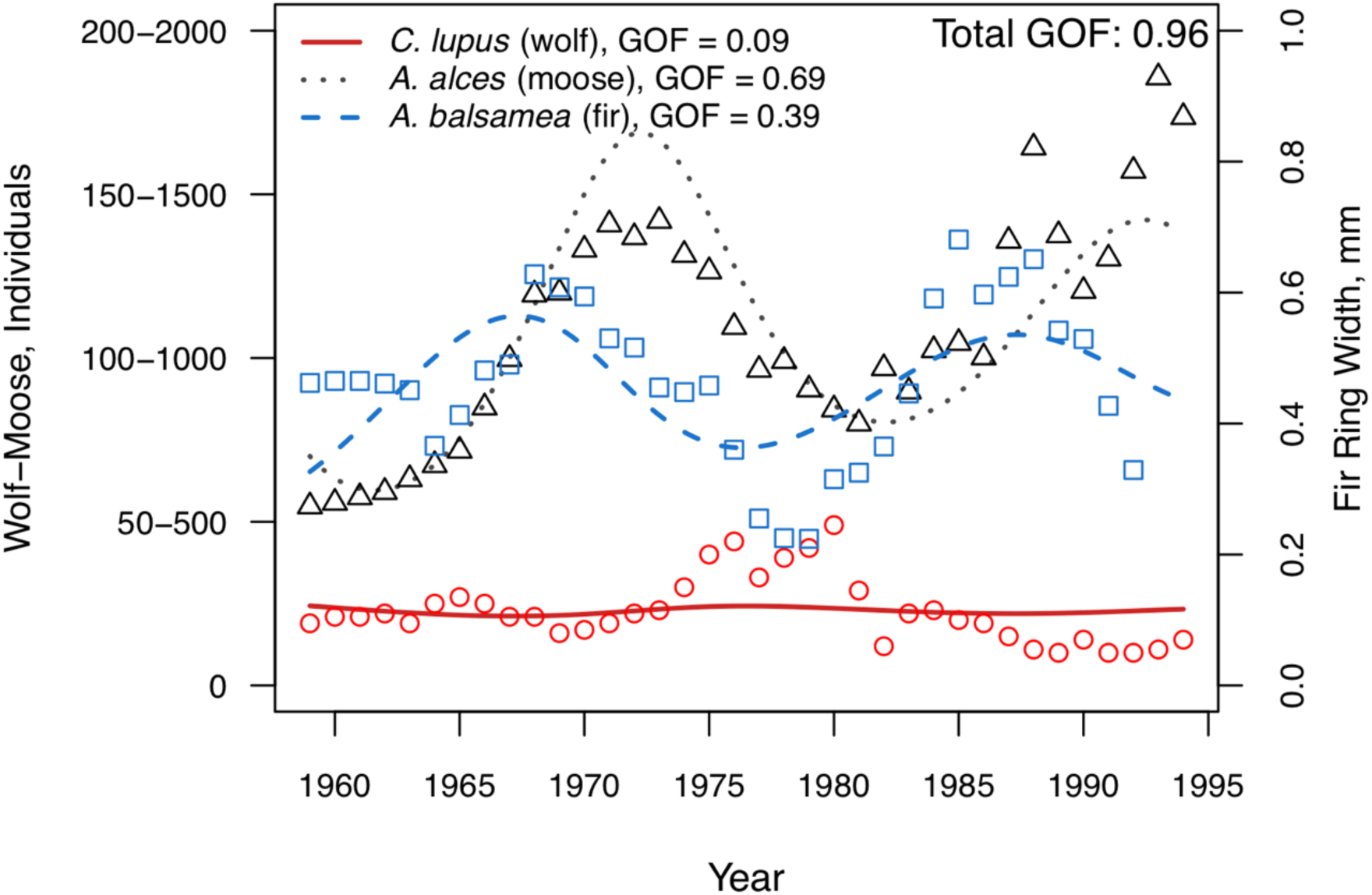
Multi-trophic dynamics for wolves, moose, and fir trees on Isle Royale from 1960 to 1994, from McLaren & Peterson (1994). Points show observations, and lines show growth curves fitted using the `lv_optim()` function. The left axis shows wolf and moose abundances, separated by a ten-fold scaling difference for easier visualization, whereas the right axis shows tree growth increments. Two methods for testing goodness of fit are shown. In the legend, values show univariate tests for each species, whereas “Total GOF” shows fit when considered across all observations simultaneously. Note that these two methods of comparison can lead to very different conclusions. See Table 1d for parameter values.

Lastly, for the Huffaker mite data, we were not able to fit the general Lotka-Volterra model to match observed dynamics (Fig. 6; Table 2e). In this system, dynamics follow sustained oscillations, but with periods and amplitudes that vary over time. However, our fitted models could only achieve either sustained oscillations of a fixed period and amplitude (top panel), or damped oscillations where the amplitude declined over time (bottom panel), but not both.

**Figure 6:**
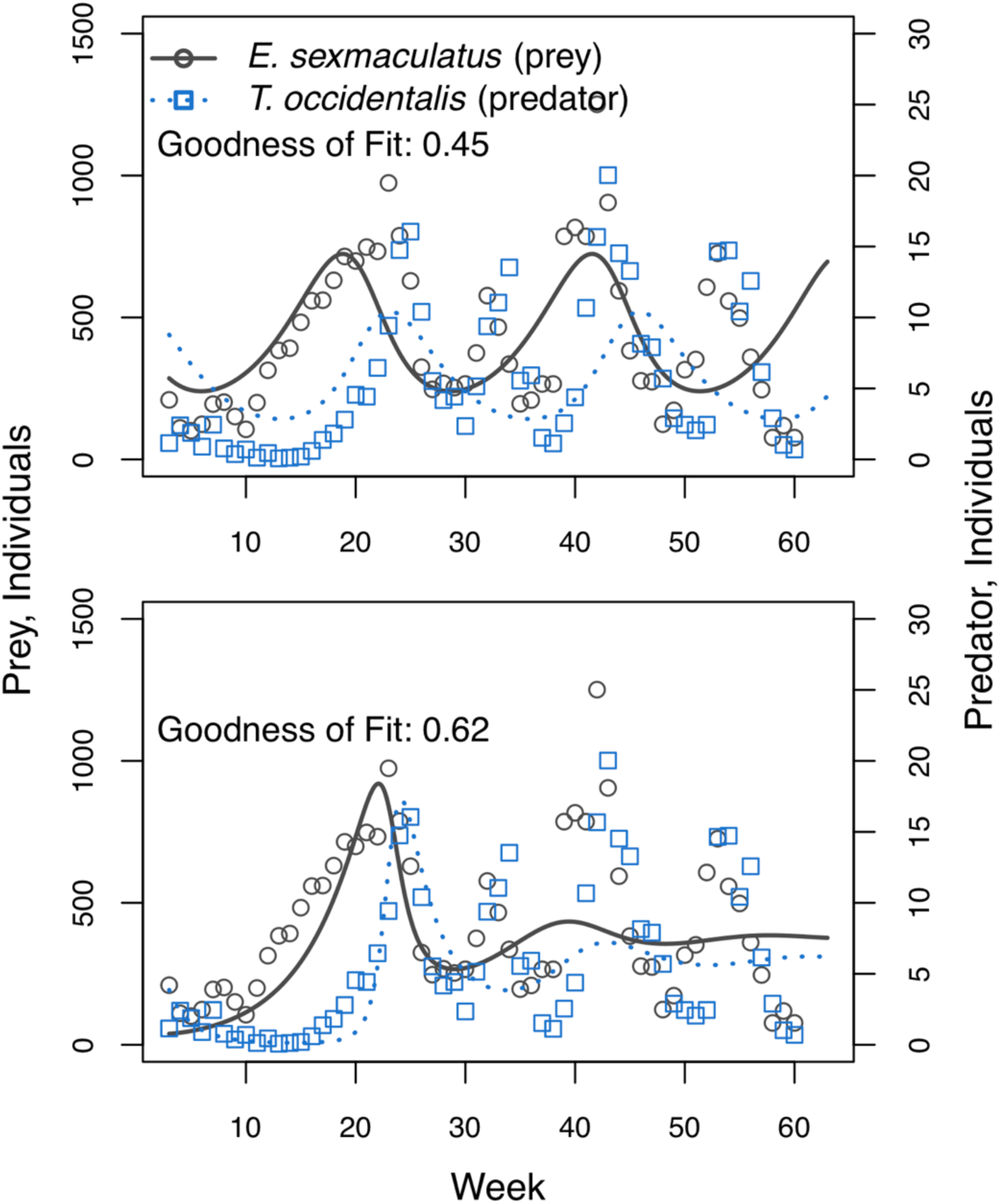
Predator-prey interactions between *Eotetranychus sexmaculatus* and *Typhlodromus occidentalis* in a spatially structured experiment carried out on a grid of oranges over 60 weeks, from Huffaker *et al.* (1963). Points show observations, and lines show growth curves fitted using the `lv_optim()` function. Top panel shows dynamics for a system where neither species directly inhibits its own growth (i.e. *a*_*ii*_ = *a*_*jj*_ = 0), whereas bottom panel is for a system where only the predator directly inhibits its own growth. Note that neither option is able to fully capture the realized dynamics, and that goodness of fit is relatively low in both panels. See Table 1e for parameter values.

## Discussion

The `gauseR` package makes it possible to analyse time series data in single-or multi-species systems using the classic Lotka-Volterra interaction models in an easy and automated way. For almost all of the examples that we consider here, models were able to accurately capture observed dynamics, and analyses could be conducted using just the default `gause_wrapper` function without any modification of the code or additional inputs from the user. Furthermore, fitted parameter values matched those expected for logistic growth, competition and predator-prey dynamics, as predicted by Lotka and Volterra. We are therefore hopeful that this combination of fitting tools, along with the package’s repository for Gause’s data, will be especially useful for teaching Lotka-Volterra’s basic equations and introducing Gause’s experimental work.

### Interpreting the models

In the monoculture experiments, our results demonstrate that *P. caudatum* followed logistic growth. As expected by theory, the species was able to increase from low abundance (i.e. *r*_*i*_ > 0), limited its own growth as its abundance increased (i.e. *a*_*ii*_ < 0), and approached a fixed carrying capacity (*K*) over time (Lotka 1925; Volterra 1926; Gause 1934a,b). For competition between *P. caudatum* and *P. aurelia*, fitted parameters indicated self-limitation by both species, and inhibition of each species by the other (i.e. *a*_*ii*_, *a*_*jj*_, *a*_*ij*_ and *a*_*ji*_ < 0). Again, these values match theoretical expectations from Lotka and Volterra, and accord with Gause’s original analyses of the data.

For the system with *D. nasutum* and *P. caudatum*, fitted models predicted that prey abundance increased in the absence of predators (i.e. *r*_*i*_ > 0), and that the predators declined exponentially in the absence of prey (i.e. *r*_*i*_ < 0,). Similarly, parameter estimates indicated that predators had a negative effect on prey, and prey had a positive effect on predators (i.e. *a*_*ij*_ < 0 and *a*_*ji*_ > 0, respectively). Jointly, these parameter values match biological expectations - i.e. that predators consume prey, and starve when prey are absent. However, in contrast to the classic formulation of the Lotka-Volterra predator-prey model, we also identified self-limitation by both species (i.e. *a*_*ii*_ < 0 and *a*_*jj*_ < 0). These parameter values resulted in damped oscillations in species abundances, as predicted by theory (Lehman *et al.* 2000). Results were similar for the wolf-moose-fir tree example, for which fitted parameters indicated self-limitation by wolves and fir trees, positive effects of moose on wolves and of fir trees on moose, and negative effects of wolves on moose and of moose on fir trees. Again, predictions from the model closely matched observations, supporting the hypothesis that top-down control of moose by wolves could explain dynamics in this system (McLaren & Peterson 1994).

In contrast, none of the models that we fitted were able to match observations from the Huffaker *et al.* mite experiments. Importantly, this limitation is a general characteristic of the Lotka-Volterra models, and not a peculiarity of our fitting methods (Lotka 1925; Hatton *et al.* 2015; Lehman *et al.* 2000). When self-limitation is absent for both species, Lotka-Volterra predator-prey models display “neutrally stable” oscillations, meaning that oscillations maintain a fixed amplitude and period. When self-limitation exists for either the prey species or the predator, or both, then oscillations become damped, meaning that over time they decrease in amplitude and period until they reach stable equilibrium values. Because the Huffaker data include persistent oscillations for which the amplitude and period vary over time, they cannot be matched by any parameterization of the Lotka-Volterra equations. Potentially, these complex dynamics are a result of the spatial structure in the Huffaker experiment, which often cannot be described with simple equations (Leibold & Chase 2018).

### Notes on optimization

For the predator-prey examples that we analyse, our findings show that parameter estimates generated from the linearized growth rates in Eq. (1b) failed to capture observed dynamics. This result accords with many other studies, which show that model performance is typically much better when parameters are tuned to match fully simulated dynamics, i.e. as is accomplished with the `lv_optim` function (Carrara *et al.* 2015; Rosenbaum & Rall 2018; Maynard *et al.* 2019). In general, the difference in fit between these two approaches is driven by the sensitivity of the Lotka-Volterra equations to noisy parameter estimates - even very small changes in parameter values can lead to large changes in model predictions (Clark & Neuhauser 2018; Maynard *et al.* 2019). We therefore suggest that in most cases, the `gause_wrapper` function, which automatically applies the optimization routine to simulated dynamics, will provide the best fits to empirical data.

Importantly, the `gause_wrapper` function applies several optimization “tricks” that help improve model performance. First, comparisons between observations and predictions are carried out on standardized abundances, such that the mean for each species is equal to one. This prevents abundant species from dominating the fitting process. For example, for the McLaren & Peterson data, optimization without standardization would closely fit dynamics of moose, the most common species, at the cost of model performance for wolves and fir trees. Second, by default, dynamics are simulated for log-transformed abundances, and are then back-transformed for comparisons to observations. This approach helps prevent computational errors in the integration from erroneously driving abundances negative. Third, we also log-transform parameter values during the optimization process, after determining their sign *a priori* from the linearized functions. This makes it possible to limit the number of positive interaction coefficients, which otherwise can cause the system to generate unrealistically high abundance values (e.g. via run-away mutualism) and crash the optimizer (Lehman *et al.* 2020). A trade-off that comes with this added stability is that this method does not allow for significance testing, because it constrains the sign of parameters. Thus, the standard errors provided by the `gause_wrapper` should only be thought of as rough approximations of the degree of uncertainty for each parameter estimate. Additionally, note that because our approach does not update predictions based on observations in each time step, it effectively assumes that all deviations between observations and predictions are due to observation error (i.e. not process noise) (Rosenbaum & Rall 2018). Users who wish to apply more sophisticated optimization methods should see the documentation and examples for the `ode_prediction` function.

There are several potential sources of observation error in the data that we analyze here. Most obviously, data digitized from figures, and especially from hand-made figures such as those of Gause and Huffaker *et al.*, are necessarily imperfect. More generally, there is reason to believe that some of Gause’s original data and figures contained some errors. In particular, in a few cases where data are available in both figures and table form, there are small differences in reported values. Similarly, many of the units reported by Gause, such as “Volume” or “Amount”, are somewhat difficult to interpret and compare across figures and experiments. Lastly, in some cases, data varies between being reported for individual replicates, to mean values summed across replicates. These circumstances present a challenge in the analysis, as they make it difficult to compare values across the entire time series, and exclude information about potentially meaningful replicate-level dynamics and between-replicate variation. Wherever possible, we have noted these inconsistencies in the metadata.

### Future directions

Given the strong correspondence between the models of Lotka (1920, 1925) and Volterra (1926) and the experimental data of Gause (1934a,b), it is perhaps not surprising that these works have had such an enormous impact on ecology over the past century. Together, these concepts have helped ecologists better describe and model their systems, and have greatly advanced conceptual understanding of how different arrangements of species and interaction types are likely to play out. Nevertheless, as the type of systems that ecologists study grow ever more complex, it seems likely that we may need to move beyond these basic models to new frameworks that allow more nuanced and dynamic interaction types (Letten & Stouffer 2019). We therefore hope that the `gauseR` package is useful in better exploring both the classic insights, and operational limitations, of Lotka, Volterra, and Gause’s framework.

## Acknowledgements

The authors declare no conflicts or specific funding sources for the work described in this manuscript.

## Tables

**Table S1:** Summary table of the datasets in the `gauseR` package. Column “name” is the name of the dataset, and can be used to load it. For all other column metadata, see the help files for each dataset in the package (i.e. via the `?name` command). See gauseR_TableS1_dataset_summarytable.csv file for contents of table. Information that is included in each dataset is marked with “Y”.

